# Genome wide estimates of mutation rates and spectrum in *Schizosaccharomyces pombe* indicate CpG sites are highly mutagenic despite the absence of DNA methylation

**DOI:** 10.1101/025601

**Authors:** Megan G. Behringer, David W. Hall

## Abstract

We accumulated mutations for 1952 generations in 79 initially identical, haploid lines of the fission yeast *Schizosaccharomyces pombe a*nd then performed whole-genome sequencing to determine the mutation rates and spectrum. We captured 696 spontaneous mutations across the 79 mutation accumulation lines. We compared the mutation spectrum and rate to another model ascomycetous yeast, the budding yeast *Saccharomyces cerevisiae*. While the two organisms are approximately 600 million years diverged from each other, they share similar life histories, genome size and genomic G/C content. We found that *Sc. pombe* and *S. cerevisiae* have similar mutation rates, contrary to what was expected given *Sc. pombe*’s smaller reported effective population size. *Sc. pombe*’s also exhibits a strong insertion bias in comparison to *S. cerevisiae*,. Intriguingly, we observed an increased mutation rate at cytosine nucleotides, specifically CpG nucleotides, which is also seen in *S. cerevisiae*. However, the absence of methylation in *Sc. pombe* and the pattern of mutation at these sites, primarily C→ A as opposed to C→T, strongly suggest that the increased mutation rate is not caused by deamination of methylated cytosines. This result implies that the high mutability of CpG dinucleotides in other species may be caused in part by an additional mechanism than methylation.

## INTRODUCTION

Spontaneous mutation is the fuel for evolution and the ultimate source of all genetic differences within and between species. A complete understanding of the mutational process, both in terms of the rate at which different mutations arise, including kind and location, and the determination of their fitness effects, is thus critical to understanding genetic variation at all levels. However, elucidating the important parameters of mutation is difficult for two reasons. First, spontaneous mutations occur infrequently, making it difficult to obtain large samples of spontaneous mutations to robustly detect patterns. One workaround is to artificially increase the mutation rate using chemical mutagens and X-rays (MULLER 1930; GREENE *et al.* 2003; BLUMENSTIEL *et al.* 2009), or genetic methods such as repair pathway knock-outs (HOFFMAN *et al.* 2004; DENVER *et al.* 2006). However, it is clear that such manipulations bias the mutational spectrum in various ways (KOORNNEEFF *et al.* 1982; GREENE *et al.* 2003). Another way to deal with the rarity of spontaneous mutations is to examine genetic differences between individuals, populations or species that have arisen via mutation. Unfortunately, because many mutations are acted upon by natural selection, this raises the second main difficulty with studying spontaneous mutations: the observed genetic variation has been acted upon by selection and is thus a biased sample of spontaneous mutations (NACHMAN and CROWELL 2000; HO *et al.* 2005). Exacerbating this problem is the finding that sites in the genome previously thought to be essentially free of selection, such as those in intronic regions, intergenic regions and four-fold redundant codon positions are, in fact, often constrained by selection, making the study of mutation at these sites biased by selection (ANDOLFATTO 2005; HERSHBERG and PETROV 2008).

One approach that has been employed to overcome the problems of rarity and selection in studying spontaneous mutations is the mutation accumulation (MA) experiment. MA experiments maintain multiple, initially identical lines at very low effective population sizes for many generations (HALLIGAN and KEIGHTLEY 2009). Lines accumulate spontaneous mutations at rates proportional to their occurrence since selection is ineffective at enriching for beneficial or decreasing deleterious mutations, unless their fitness effects are large (SUNG *et al.* 2012a). Each line accumulates only a few mutations but, by having numerous lines, several hundred mutations can be captured across the lines. Whole-genome sequencing (WGS) of the MA lines allows mutations to be identified and used to estimate the frequency and spectrum of spontaneous mutation. This approach has been used to examine spontaneous mutations in MA lines in a number of eukaryotic species including *Arabidopsis thaliana, Caenorhabditis elegans, Chlamydomonas reinhardtii, Dictyostelium discoideum, Drosophila melanogaster, Paramecium tetraurelia*, and *Saccharomyces cerevisiae* (KEIGHTLEY *et al.* 2009; DENVER *et al.* 2012; NESS *et al.* 2012; RUTTER *et al.* 2012; SAXER *et al.* 2012; SUNG *et al.* 2012b; ZHU *et al.* 2014). The largest study of this kind to date, in *S. cerevisiae*, yielded a total of 924 spontaneous mutations (ZHU *et al.* 2014).

The species that have been utilized in WGS MA experiments are a somewhat miscellaneous group, notable for their ease of culture and rapid generation times. While we have learned a great deal by comparing mutation rates and spectrum estimates across them (LYNCH 2010), it is difficult to generate and test hypotheses for differences in the observed spectra across such a disparate group of species. To begin to remedy this problem, we chose to perform a MA experiment in the haploid, fission yeast *Schizosaccharomyces pombe* to specifically compare results to *S. cerevisiae*. Originally isolated from millet beer, *Sc. pombe* is an important model organism in molecular biology (HUMPHREY 2000; WOOD *et al.* 2002). While *Sc. pombe* and *S. cerevisiae* are distantly related evolutionarily, with a divergence time of 600-1200 million years (HECKMAN *et al.* 2001; DOUZERY *et al.* 2004), they exhibit similar life histories as single-celled, sexual yeasts and they are cultured in the lab using essentially identical methods. In addition, they have similar genomic G/C content (36.06% in *Sc. pombe* versus 38.29% in *S. cerevisiae*) and genome size (13.8 Mb in *Sc. pombe* (Wood et al. 2002) versus 12.1 Mb in *S. cerevisiae* (Goffeau et al. 1997)). While similar, they differ in two features that are expected to have implications for their mutation rate and spectrum. *Sc. pombe* has a smaller effective population size (BROWN *et al.* 2011), and is a haploid as opposed to a diploid species in nature. The similarities and differences between these two yeast species allowed us to make five *a priori* predictions concerning mutation rate and spectrum in *Sc. pombe*.

Our first prediction was that the genome-wide single-nucleotide mutation (SNM) rate would be substantially higher in *Sc. pombe* than in *S. cerevisiae*. This prediction comes from previous mutation rate estimates in *Sc. pombe* based on reporter genes that average 8.2 × 10^-10^ single nucleotide substitutions per base per generation (summarized in Lynch 2010), which is 4.8 fold higher than the most recent estimate in *S. cerevisiae* (ZHU *et al.* 2014). Further, genome-wide nucleotide diversity (π) is similar in these two species: 0.0030 in *Sc. pombe* (JEFFARES *et al.* 2015) and 0.0057 in *S. cerevisiae* (LITI *et al.* 2009). The equilibrium level of π at a locus is expected to equal 2 *n N_e_ μ*, where *n* is the ploidy (*n* = 1 for haploids and *n* = 2 for diploids), *N*_*e*_ is the effective population size and *μ* is the per generation mutation rate at the locus. Assuming *Sc. pombe*’s mutation rate estimate is accurate, its effective population size is then predicted to be ∼2.5 times smaller than *S. cerevisiae*. Smaller effective population sizes are thought to limit the evolution of replication fidelity because the strength of natural selection favoring increased accuracy declines as accuracy increases. Eventually population error rates decline to a point where they encounter the drift barrier (SUNG *et al.* 2012a); the point at which the fitness advantage of an additional reduction in the mutation rate is the same magnitude as the strength of genetic drift, on the order of the reciprocal of 2*N*_*e*_*u*. Using the relationship reported in Sung *et. al.* (2012), a 2.5 fold reduction in *N*_e_ will result in a 2.3 fold increase in the mutation rate. Given the most recent estimate of the SNM rate in *S. cerevisiae (ZHU et al. 2014)*, we expected to estimate a rate in *Sc. pombe* that was at least 2.3 fold larger; 3.8 × 10^-10^ per base per cell division. We note, that the estimated *N*_*e*_ is based on the reporter-construct estimated *μ*, which is then used to estimate the expected genome-wide mutation rate. Thus, if the reporter-construct estimated mutation rate is inaccurate, then the expected genome-wide mutation rate will be different.

Our second prediction was that most of the SNM biases, which are the relative rates of mutations among the different base pairs, in *Sc. pombe* would be essentially the same as in *S. cerevisiae*. This prediction comes from the fact that the genomic G/C content of *Sc. pombe* (36.06%) is similar to *S. cerevisiae* (38.29%). Interestingly, the SNM biases observed in *S. cerevisiae* predict a lower G/C content (32%) than observed (Zhu et al. 2014), suggesting another force acts on G/C content. One possibility is selection. If weak selection drives G/C content to a higher equilibrium than predicted based on SNM bias in both species, then we expect *Sc. pombe* to be closer to the mutation-bias equilibrium than *S. cerevisiae*, i.e. lower, as observed, because of its smaller effective population size, which reduces the efficacy of selection. The SNM bias in *S. cerevisiae* shows an elevated mutation rate of C:G base pairs (ZHU *et al.* 2014). A major surprise concerning the elevated mutation rate at C:G base pairs was the finding that their mutagenicity is affected by trinucleotide context. Specifically when C:G is the middle base pair in CCG (equivalent to CGG) and TCG (equivalent to CGA) trinucleotides, the mutation rate is elevated even more than for other C:G base pairs. Both of these trinucleotides include a CpG dinucleotide, which is a well-characterized target for methyl-transferases (BESTOR and VERDINE 1994). The bias at CpG dinucleotides was in the C → T direction. For this reason, the finding of elevated C:G mutation at CCG and TCG trinucleotides was interpreted as indicating a very low occurrence of DNA methylation in *S. cerevisiae* (ZHU *et al.* 2014), in agreement with previous work (TANG *et al.* 2012). *Sc. pombe* is believed to lack DNA methylation so we did not expect to see an elevation of the G:C mutation rate at CpG sites (ANTEQUERA *et al.* 1984).

Our third prediction was that among small insertions and deletions (indels) there would be an insertion bias. Small indels occur primarily at microsatellites, presumably primarily due to slippage of the DNA polymerase during replication (LEVINSON and GUTMAN 1987). In *S. cerevisiae*, indels of less than 50bp were moderately biased towards deletions (18 deletions versus 8 insertions) across the diploid MA lines (ZHU *et al.* 2014). However, analysis of indels in haploid MA lines of *S. cerevisiae* showed a bias towards insertions (34 insertions and 8 deletions at microsatellite loci) (LYNCH *et al.* 2008). Since *Sc. pombe* is a natural haploid and was passaged as such in our MA experiment, we hypothesized that we would see a bias towards insertions for small indels in this species.

Our fourth prediction was that mutations resulting from non-recombinational repair of double strand breaks would be more common in haploid *Sc. pombe* than in diploid *S. cerevisiae*. Non-recombinational repair is substantially more mutagenic than recombinational repair (DALEY *et al.* 2005). In a diploid cell, there is always a non-sister homolog present that can be used for recombinational repair. However, in a haploid cell, recombinational repair is only possible during the S or G2 phase of the cell cycle, when a sister chromatid is present. Repair of double-strand breaks is thus expected to be more error-prone in a haploid because of the higher likelihood of using non-recombinational repair. While double strand breaks cannot be directly observed or always inferred with certainty in the MA framework, their occurrence can be indicated if they are inaccurately repaired by the presence of multiple mutations in close proximity (STRATHERN *et al.* 1995; HOLBECK and STRATHERN 1997). In diploid *S. cerevisiae*, three double mutations, which consist of two SNMs adjacent to one another, were observed across the MA lines (ZHU *et al.* 2014). The relative occurrence of double SNMs was thus ∼0.35% compared to SNMs. In addition, there were five “complex mutations”, in which multiple SNMs, and often small indels, were in close proximity, giving a relative occurrence of ∼0.58% compared to SNMs. If both double and complex mutations were due to error-prone double-strand break repair, then the observed number of these mutations accounted for ∼1% of SNMs. The haploid MA experiment in *S. cerevisiae* was too small to detect such rare events (LYNCH *et al.* 2008). However, our MA experiment is large enough that we predicted we would see a substantial increase in the rate of double mutations and complex mutations in *Sc. pombe* compared to *S. cerevisiae*, assuming these events are indeed caused by repair of double stranded breaks.

Our final prediction was that we would not observe aneuploidy in *Sc. pombe*. Since *Sc. pombe* is haploid, loss of a single chromosome (nullisomy) results in loss of all copies of the genes on that chromosome, which would likely be lethal and thus unobservable. In addition, gain of a single chromosome (disomy) would result in a doubling of gene dose for all genes on the chromosome, which is a larger increase than the 1.5 fold increase that occurs with trisomy in diploid *S. cerevisiae*. If deleterious effects due to gene dosage were increased with larger differences in dosage across genes, this would further reduce the likelihood of observing aneuploidy. In addition, *Sc. pombe* has only three chromosomes, implying that many more genes would be affected by an aneuploidy event than in *S. cerevisiae*, which has a similar genome size but 16 chromosomes. The only instance of aneuploidy reported in previous work in *Sc. pombe* is disomy of chromosome III, which was highly deleterious (NIWA *et al.* 2006). For these reasons, we predicted we would see no nullisomy and likely no disomy in any of our MA lines.

This study presents the first genome wide estimates of mutational parameters for *Sc. pombe*. We examined 79 MA lines, cultivated as haploids for an average of 1952 generations. We were able to identify a total of 696 mutations. These mutations allowed us to calculate precise estimates of the mutation rate and spectrum for *Sc. pombe.*

## MATERIALS AND METHODS

### Mutation Accumulation Lines

*Sc. pombe* MA lines were passaged in the same manner as described for *S. cerevisiae* (JOSEPH and HALL 2004). Briefly, the haploid ancestral line, 972 h- (ATCC 26189), was streaked onto rich, solid YPD medium (1% yeast extract, 2% peptone, 2% dextrose, 2% agar) and incubated at 30°C. From the streaked ancestor, 96 random isolated colonies were selected after 48 hours and used to found 96 MA lines. Lines were cultured six to a YPD plate and bottlenecked by randomly selecting one isolated colony per line every 48 hours and transferring to a new plate. Lines were passaged for a total of 100 transfers (200 days). Every ten transfers, a random colony from each line was frozen and stored in 15% glycerol at - 80°C.

Every 10 transfers, photographs were taken of all 96 MA lines. From these photographs, colony size was measured for five random colonies per line using ImageJ (SCHNEIDER *et al.* 2012). In addition, the numbers of cells per colony for colonies of various sizes were recorded at transfer 10 (T10) and 100 (T100) by suspending a single colony in 1ml of water and counting individual cells using a haemocytometer. These measurements were used to determine standard curves of colony size versus cell number at these two transfers. Measurements of average colony sizes at every tenth transfer were then used to calculate the average number of cells at those transfers. From the average number of cells at every tenth transfer, the number of cell generations per transfer, and across the entire experiment, was estimated, and the effective population size across the experiment determined.

### Sequencing

MA lines were cultured from frozen T100 stock on solid YPD medium at 30°C for 48h. A single colony from each line was selected, inoculated into 3mL liquid YPD, and incubated on a rotator at 30°C for 48h. Cells were then pelleted and DNA was extracted using the YeaSTAR kit (Zymo Research) protocol with chloroform and an extended digestion time with zymolase of 2.5h at 37°C. Whole genome shotgun libraries were prepared by the Georgia Genomics Facility using the Kapa Library Low-Throughput Library Preparation Kit with Standard PCR Amp Module KK8232 with dual SPRI size selection cleanup to generate 100bp paired end fragments with ∼300bp inserts. After 7 cycles of PCR the libraries (96 haploid MA lines and 2 haploid ancestor samples) were pooled across two lanes of Illumina HiSeq 2000 machines (Sequences to be uploaded to SRA).

### QC, Mapping, and Identification of Mutations

Sequence reads from each library were quality controlled with the ea-utils and fastx toolkit in order to remove low quality reads and residual adaptor sequence (GORDON and HANNON 2010; ARONESTY 2011)(Workflow deposited at https://github.com/behrimg/Scripts/blob/master/Hall_Projects/Pombe/MA_Pipeline.txt). Based on the workflow outlined in Zhu et al. (2014) and adjusted for a haploid dataset, QCed reads were then mapped to the *Sc. pombe* reference genome ASM294v2.24 with BWA v1.1.2, sorted and indexed with SAMtools v1.0, and assigned line identification numbers with Picard Tools v1.87 (WOOD *et al.* 2002; LI and DURBIN 2009). Duplicated reads were marked with Picard Tools and removed, and then the remaining sequence reads were locally realigned with GATK v3.2.2 (MCKENNA *et al.* 2010). SNM and indel variants for each line and the ancestor were identified simultaneously using GATK’s Unified Genotyper tool with parameter settings for haploid organisms. The resulting VCF files were converted to tab delimited text using VCFtools v0.1.12a vcf-to-tab function (DANECEK *et al.* 2011). All differences between the MA ancestor sequences and the reference sequence were identified to determine the sequence of the ancestor. The differences between each MA line and the reference were determined and those that were present in the ancestor were ignored. In order to call a variant, a minimum of four reads with ≥75% of the reads favoring the variant allele was needed. Regions of the genome that corresponded to centromeres, telomeres, and mating type loci (approximately 248kbp) were excluded from the analysis to avoid inaccurate mapping. This was in addition to the two tandem rDNA repeat arrays on chromosome III accounting for 1,465kbp, which are excluded from the reference genome. Identified SNMs and small indels were annotated using Ensembl’s variant effect predictor (VEP) while flanking regions were determined using the fill-fs program from the VCFtools package (LI and DURBIN 2009; MCLAREN *et al.* 2010).

Presence of medium and large structural variants were investigated using the Delly software package (RAUSCH *et al.* 2012) and variants that passed Delly’s QC were investigated further using the integrated genome viewer (IGV) v2.1.23 (ROBINSON *et al.* 2011). When IGV supported a structural variant call, the variant was tested with PCR (Primers in Supplementary Materials).

Sequencing also allowed the detection of across-line and other microbial contamination. Across-line contamination was deemed to have occurred if any two lines shared an identical new mutation. When this happened, one of the lines (chosen by coin flip) was discarded from the remainder of the analysis.

### Gene Expression

To estimate gene expression levels for our ancestor strain, we sequenced mRNA from 10 biological replicates. We selected 10 colonies, inoculated each into 3mL liquid YPD medium, and incubated on a rotator at 30°C for 48h. After 48h, mRNA was extracted using the MasterPure Yeast RNA Purification kit (Epicentere). mRNA libraries were constructed using the Illumina Truseq mRNA Stranded Kit, amplified using 13 cycles of PCR and sequenced on an Illumina HiSeq 2500. Libraries were sequenced as 100bp single-end reads (Sequences to be uploaded to SRA). Sequenced reads were QCed in the same manner as genomic sequencing reads, reference-mapped with TopHat v.2.0.13 (TRAPNELL *et al.* 2009), and assembled with Cufflinks (TRAPNELL *et al.* 2010). The log-median FPKM for each site across ancestor replicates was chosen to represent the level of expression at that site.

### Verification of Identified Mutations

Five lines were randomly selected to verify the mutations that were identified bioinformatically with Sanger sequencing (primer sequences listed in Supplementary Materials). Primers were designed using Primer3 (ROZEN and SKALETSKY 1999) and PCR products destined for sequencing were cleaned using a standard Exo-SAP protocol (BELL 2008), and sequenced with an ABI BigDye Terminator Cycle Sequencing Kit (Applied Biosystems, Foster City, CA). Completed sequencing reactions were submitted to the Georgia Genomics Facility and analyzed using an Applied Biosystems 3730xl 96-capillary DNA Analyzer. Additionally, false negative rates for SNMs and small indels were determined by determining the number of mutations present in the ancestor, which were not identified bioinformatically across the 79 MA lines.

### Relative mutation rate analysis

To investigate the effects of G/C content, replication time, and transcription rate, relative mutation rates were calculated. G/C content was determined in non-overlapping 10 kb windows across the entire genome. The number of SNMs was determined for each window and separated into 14 bins representing G/C content rounded down to the nearest 1%. Relative mutation rates were determined for each bin by determining the per base pair mutation rate for each bin and dividing by the observed genome-wide per base pair mutation rate. Bins were then separated into groups so that each group contained roughly equal numbers of SNMs but not necessarily equal numbers of bins. Weighted averages and standard errors for each group were determined by weighting each bin by the amount of the genome covered by the bin. The same bins were used to determine relative mutation rates at A/T and G/C bases by calculating the per base pair mutation rate for just A/T and G/C bases.

Replication time was determined genome wide by assigning each SNM a replication time based on the closest origin of replication (HEICHINGER *et al.* 2006). SNMs were separated into 17 bins, each representing one minute of S phase of the cell cycle. Relative mutation rates were then determined in an analogous manner as for G/C content.

Transcription rate was determined genome wide and transcripts were separated into 15 bins based on gene expression increasing from 0 to 3.5 by 0.25 log_10_FPKM. SNMs were assigned a bin based on the transcript in which they reside, SNMs not mapping to a transcript were assigned a gene expression of 0. Relative mutation rate was then determined in an analogous manner as G/C content and replication time.

## RESULTS

### MA Lines

The relationship between colony area and the number of cells in the colony did not differ between transfers 10 and 100 (data not shown). The regression between colony size and transfer number indicated that average colony size declined over the course of the experiment such that the number of cells at T100 was approximately half the number at T10, indicating an average reduction in per-generation growth rate of approximately 5%. In diploid *S. cerevisiae*, a trend towards reduced colony size was not observed (Joseph and Hall 2004), presumably because new mutations accumulated in the heterozygous state and thus had a smaller effect on fitness due to masking. The average number of cells in a colony was used to estimate the average effective population size in a MA line as 10.26 cells. This small effective population size implies that deleterious (beneficial) mutations with a fitness effect greater than ∼0.097 would be underrepresented (overrepresented) due to the action of selection (HALL *et al.* 2008; LYNCH *et al.* 2008). Mutations with smaller fitness effects are expected to accumulate approximately at random in the MA lines.

Mapping of sequencing reads to the *Sc. pombe* reference genome revealed that coverage was uniform across all three chromosomes except for telomeric regions in those lines with average sequencing depth below 43x, where telomeric sequences appeared to be affected by amplification bias (Figure S1). After sequencing and identification of nucleotide variants, we discarded seventeen of the 96 MA lines due either to contamination by a different microbial species (6 lines) or to across MA line contamination (11 lines), which left 79 lines for further analysis. We note that for most of the contaminated lines (10 of 11), all nucleotide variants were shared indicating that contamination likely occurred either at one of the last transfers, or when the lines were removed from the freezer for DNA sequencing.

Approximately 248kbp, which is ∼2% of the sequenced genome, was excluded from the analysis to ensure precise identification of new mutations. This included the low-complexity sequence, comprising centromeres, telomeres, the mating type region, and the representative rDNA repeat belonging to the two tandem rDNA repeat arrays on chromosome III, which had been previously excluded from the 12.47 Mb reference genome (WOOD *et al.* 2011).

### Differences between the MA ancestor and *Sc. pombe* reference genome

Sequencing of the ancestor revealed a total of 272 differences between it and the reference genome. These differences included 85 single nucleotide mutations (SNMs), 129 small insertions (< 50 bp), 47 small deletions (< 50 bp), 6 double SNMs and 5 complex mutations (Figure S2). We define double SNMs as two SNMs that occur within 50 nucleotides of one another. Complex mutations are three or more SNMs and/or indels occurring within 50 nucleotides of one another. Both complex mutations and double SNMs were deemed to be non-independent events and were thus analyzed separately. Of the 85 SNMs, 41 were transitions and 44 were transversions, giving a transition to transversion ratio of 0.93. Within transitions, after correcting for genomic G/C content and assuming that all changes were mutations from the reference strain sequence to the ancestral strain sequence, G:C → A:T or T:A mutations were 1.38 times more frequent than A:T → G:C or C:G mutations (Figure 1).

**Figure 1.**
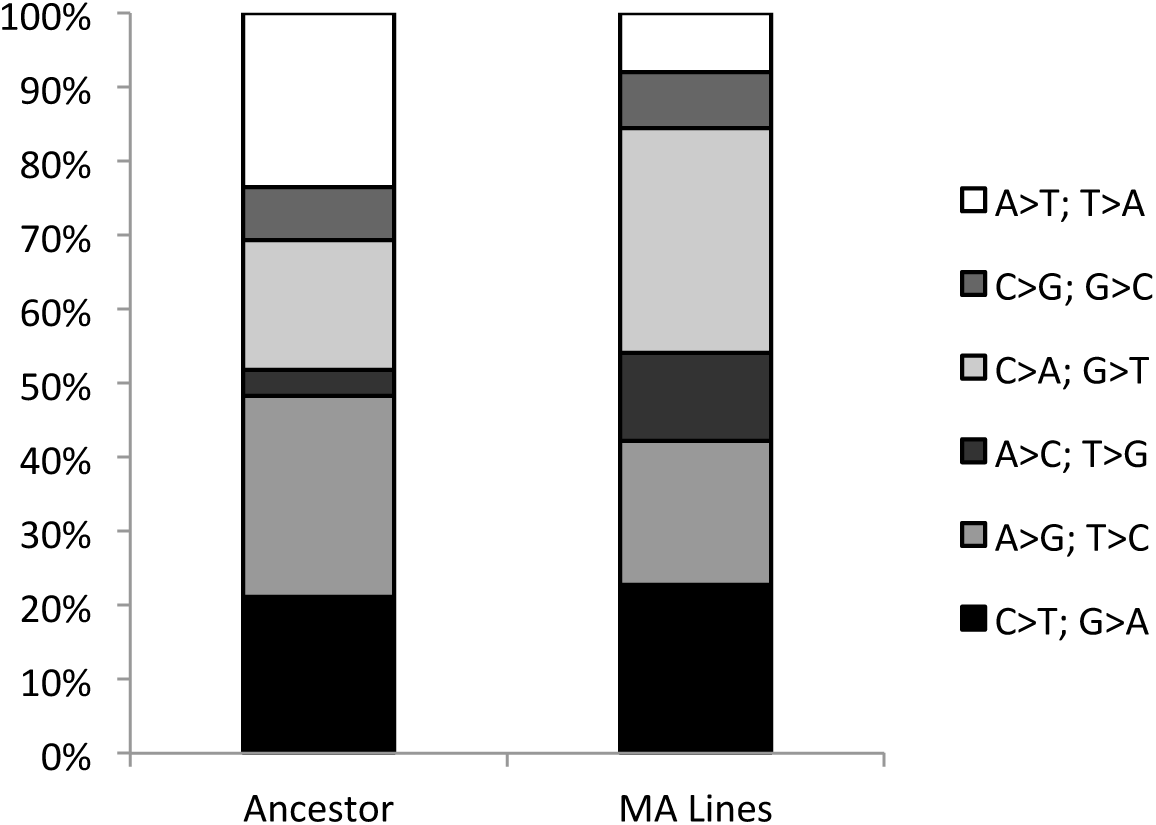
Summary of mutations for each of six possible nucleotide changes for both the ancestor and the MA lines compared to the reference genome.

The distribution of SNMs across chromosomes was not significantly different from the expectation based on chromosome length (X^2^ = 5.45, p= 0.06). This result held whether we tested all three chromosomes, or just chromosomes I and II (X^2^ = 3.05, p = 0.06), which together represent 81% of the genome. The distribution of small insertions was non-random, with chromosome I (5.58 Mb) containing 75, which is greater than the 57 expected based on chromosome length (X^2^ = 12.21, p = 0.002). Other mutations, including small deletions, complex mutations, and double SNMs, did not show bias across chromosomes, after accounting for length (Figure S3).

### Differences between the MA lines and the MA ancestor

#### Single Nucleotide Mutations

Across the 79 MA lines, 326 SNMs arose during MA, which gives the single-nucleotide mutation rate (± 1 standard error) for *Sc. pombe* as 0.170 ± 0.013 × 10^-9^ per base per generation (Figure 2). The standard error of the estimate is based on the variance across MA lines. This mutation rate estimate does not include the 123 SNMs that were in double or complex mutations. Including SNMs in double and complex mutations increases the genome-wide mutation rate estimate to 0.234 ± 0.023 × 10^-9^ per base per generation.

**Figure 2.**
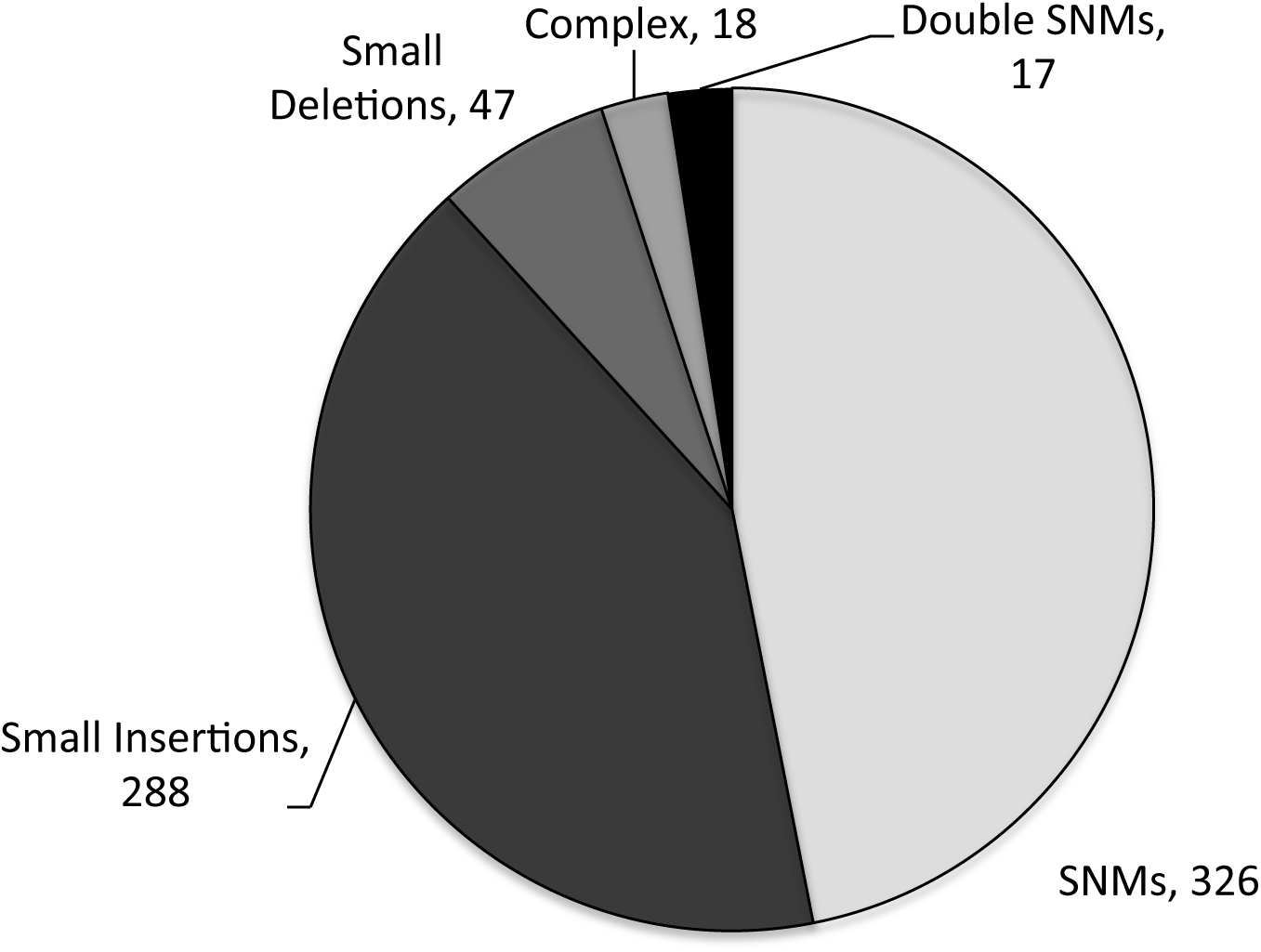
Summary of types of mutations identified across 79 MA lines.

The number of SNMs varied across the 79 MA lines from zero (eight lines) to 13 (one line) with an average of 4.13 SNMs per line (ignoring doubles and complex mutations). Surprisingly, the distribution of mutations across MA lines was not consistent with a Poisson distribution (χ^2^; p < 0.001). A negative binomial distribution (gamma-Poisson), with mean = 4.13 and overdispersion parameter = 2.06, could not be rejected (χ^2^; p = 0.954, Figure 3). Using Akaike’s information criterion (AIC) the negative binomial was a substantially better fit to our data than the Poisson (Poisson AIC = 15.58, Negative Binomial AIC = 4.45; Poisson is 0.0034 as likely to explain the data). SNMs occurred at random with respect to protein encoding genes: 52.4% of SNMs were in the 57% of the genome that is protein coding (Fisher’s exact, p = 0.27), and 3.6% of SNMs were in the 3% of the genome that is intronic sequence (Fisher’s exact, p = 0.83), suggesting that selection was inefficient during MA.

**Figure 3.**
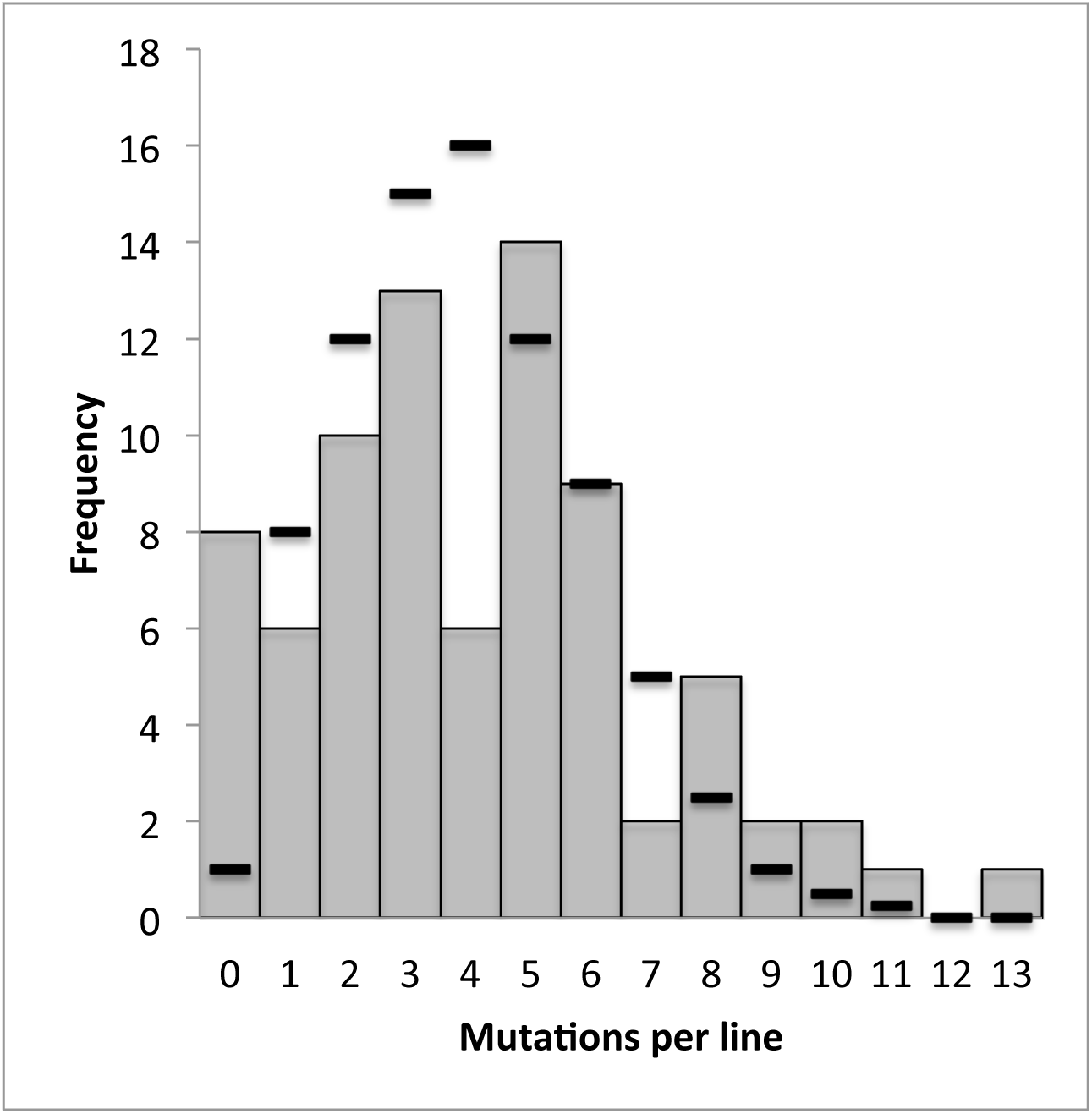
Histogram of SNM numbers per line across the 79 MA lines. The distribution is consistent with a negative binomial (P = 0.95, χ^2^ test). Dashes represent the expected numbers for the best-fit negative binomial distribution (mean = 4.13, overdispersion = 2.06). The total number of SNMs across all 79 MA lines is 326.

The distribution of SNMs across the three chromosomes was borderline significantly different from the expectation based on chromosome length (X^2^ = 4.99, p= 0.08)(Figure S3). The distributions of all other mutation classes (small insertions, deletions, complex, double SNMs) were not significantly different than expected based on chromosome length.

#### Single Nucleotide Mutation Biases

Ignoring SNMs in complex and double mutations, we observed a transition to transversion (Ts/Tv) bias of 0.72 (Figure 1). Within transitions, G:C → A:T mutations were 2.02 times more frequent than A:T → G:C. Within transversions, G:C → T:A mutations were 4.55 times more frequent than A:T → C:G. Across transitions and transversions, G/C → A/T mutations were 2.97 times more frequent than A/T → G/C mutations. If the mutational process were the sole determinant of the G/C content in *Sc. pombe*, the equilibrium genome G/C content would be 25.14%, far less than the 36.06% observed in the reference genome.

Local G/C content did not affect mutation rate. We determined the G/C content in 10kb windows across the genome and then separated the data into groups such that each contained a similar number of SNMs. Regardless of whether we split the data into two or three groups, we found no significant difference in mutation rate across them, suggesting that local G/C content has no effect on local mutation rate. Data for the two-group case where the division is for a GC content of 36% is shown in the Supplemental Materials (Figure S4A).

Replication time also did not affect mutation rate. We assigned each nucleotide in the genome a replication time based on the time at which the closest origin of replication (ori) initiates replication during mitosis (HEICHINGER *et al.* 2006). There is little variation in the replication initiation times of different oris in *Sc. pombe*, with all oris firing between 68 and 85 minutes after release from G2 arrest (HEICHINGER *et al.* 2006). Further, published initiation times are reported in 1-minute intervals and so we could not split times into smaller intervals. We separated replication times into three groups and found no significant differences in mutation rate for different replication times (Figure S4B).

Expression level was not associated with mutation rate. Using RNA-seq data collected from our ancestor line, we examined whether gene expression, measured as transcript abundance, influenced local mutation rate. Transcription rates were split into 15 bins that were then separated into two groups. These two groups did not differ in mutation rate.

Since our protocol for creating RNA libraries employs poly-A capture, tRNAs, which likely have very high expression levels (WOOD *et al.* 2011), are not represented in our transcriptome. When SNMs that occur in tRNA are not included, there is no effect of expression level on mutation rate, as described in the previous paragraph. However, when we include tRNAs and the associated SNMs, and assume that they are in the highest expression category (WOOD *et al.* 2011), we find a nonsignificant trend for higher relative mutation rate in highly expressed genes (Figure S5). This difference is not statistically significant because the relative mutation rate for the bin containing the highest expressed genes has a very large standard error.

We did find an effect of trinucleotide context, which is the identity of the nucleotides immediately before and after the mutated nucleotide, on mutation rate (ZHU *et al.* 2014). Each nucleotide position within the genome, along with its neighboring bases, was assigned to one of the 64 trinucleotide possibilities. Ignoring strand orientation allowed us to reduce the 64 possibilities to 32 groups, defined such that complementary trinucleotides belong to the same group (e.g. GCA and TGC are in the same group). Mutation rates at the center position across the 32 groups were quite variable (Figure 4). We analyzed the 16 groups with A (or T) at the center position separately from those with G (or C) because C:G base pairs have a higher mutation rate than A:T base pairs, as is clearly apparent in Figure 4. After correcting for multiple comparisons, we found context did not affect mutation rate at A:T base pairs. In contrast, at C:G base pairs, mutation rates were increased in the CCG (equivalent to CGG), GCG (equivalent to CGC), and GCA (equivalent to TGC) trinucleotide groups (t-test; p = 0.0003, p = 0.0001, and p = 0.0009 respectively; Bonferroni corrected critical p-value = 0.0031). Interestingly, two of these trinucelotides are two of the four trinucleotide groups that contain a CpG dinucleotide, where the C is the at the center position. If increased CpG mutation rate was due to deamination of methylated cytosine (giving thymine), then we would expect CpGs to mutate to TpGs. However, of the 44 mutations occurring at CpG sites, 23 were C → A mutations (52.3%), 18 were C → T (40.1%) and 3 were C → G (6.6%); which is no different than expected based on *Sc. pombe*’s mutational spectrum (X^2^ = 1.76, p = 0.41). Additionally, for the GCA trinucleotide group, C → A mutations were over-represented (X^2^ = 8.91, p = 0.011).

**Figure 4.**
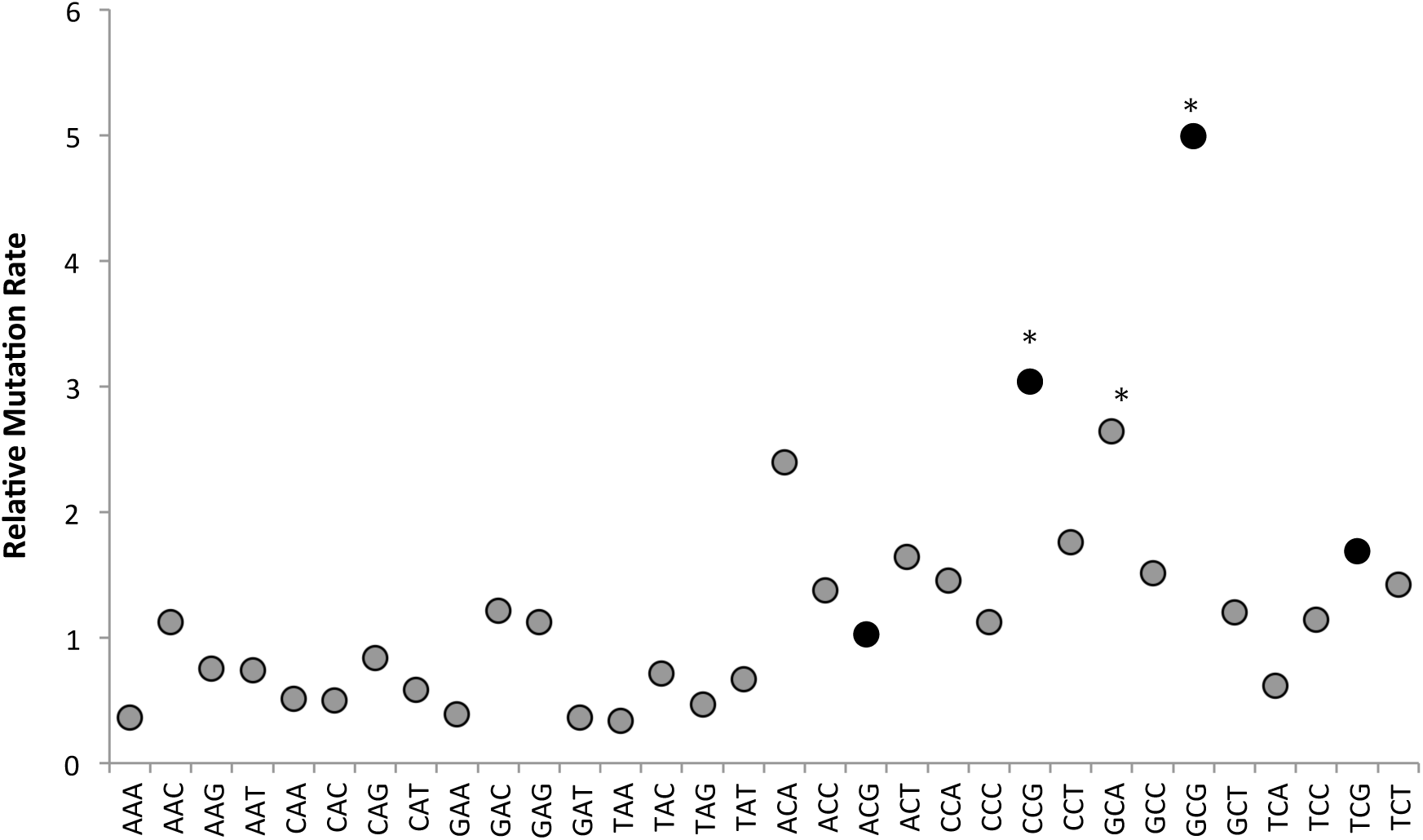
Mutation rate is affected by context. Relative single nucleotide mutation rate adjusted by its trinucleotide context. Trinucleotide classes represent mutation rates of both strand orientations. For example aCa trinucleotide class includes overall mutation rate at aCa and tGt sites. Mutation rate is shown relative to the average single nucleotide mutation rate across all sites (= 1.7 × 10-^10^ per base, per generation). The average, relative mutation rate of 1.8 at G:C bases shows clear overall elevation over the corresponding rate of 0.66 at A:T bases. The four black-filled points are those trinucleotides with a C in the central position with a G as the 3’ neighbor. * p ≤ 0.003 Bonferroni corrected t-test.

#### Small Insertion and Deletion Mutations

Across the MA lines, we identified 335 small indels of less than 50 bp, including 288 insertions and 47 deletions. A complete list of indels is in the Supplemental Materials. Average insertion size was 1.6 bp while average deletion size was 3.1 bp, with insertions occurring 6 times more frequently than deletions, resulting in a net gain in DNA sequence across all lines of 340 bp. Small indels occurred as frequently as SNMs across the MA lines and the resulting spontaneous indel rate, 0.174 × 10^-9^ indels/base/generation, is essentially identical to the mutation rate calculated for SNMs, ignoring double SNMs and complex mutations. Indels however were not randomly distributed with respect to genomic features. They were substantially under represented in protein coding sequence (observed: 33, expected: 191; Fisher’s Exact Test: p < 0.001), occurred as frequently as expected in introns (observed 21, expected 10; p = 0.064), and were over represented in non-coding regions (observed 288, expected 134; p < 0.001).

#### Effects of SNMs and Indels

We annotated the expected functional effects of our SNM and small indel mutations using Ensembl’s variant effect predictor (VEP) software (MCLAREN *et al.* 2010). For the 183 SNMs that occurred within coding regions, 53 were synonymous, 113 were missense, 12 were within introns, 1 was within a splice donor site, 1 was within a splice acceptor site, 2 were nonsense and 1 changed a termination codon (UAA) into another termination codon (UAG) (Figure 5). The ratio of synonymous to missense changes (0.398) was higher but not statistically different from the expected ratio (0.323), computed using randomly generated protein coding sequences (Z-test for proportions: 1.104, p = 0.271) (Figure S6) (GRAUR 2003). This suggests selection did not substantially reduce the number of missense mutations captured during MA.

**Figure 5.**
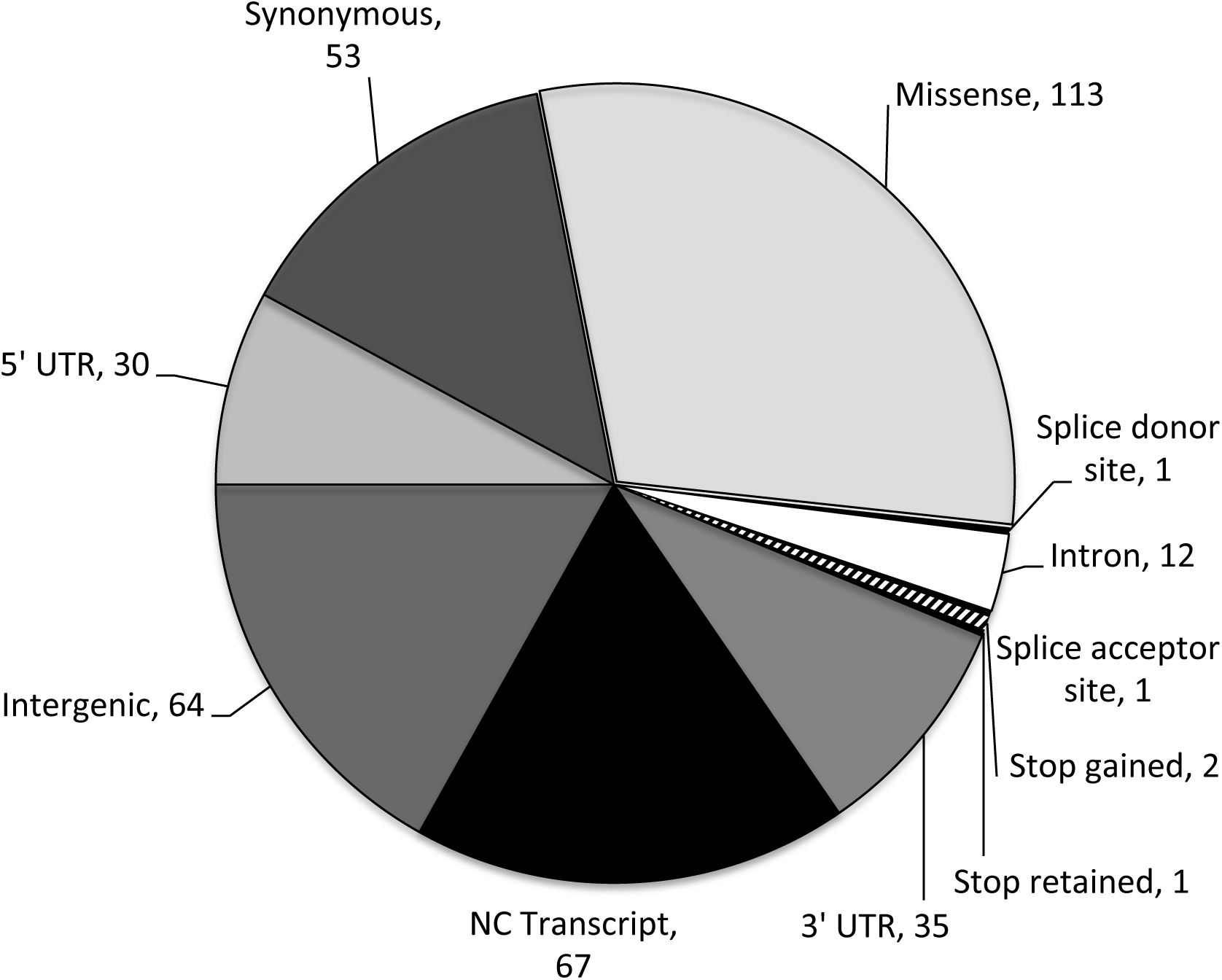
Summary of the locations and predicted functional consequences of the 326 accumulated SNMs. Numbers of SNMs observed across all 79 MA lines are shown for each category. Sum of numbers (379) is greater than the observed number of mutations across all MA lines (326) because some mutations had multiple predicted functional effects. NC = non-coding. UTR = transcribed, untranslated region.

Among the 54 small indels that occurred within protein and intronic coding sequence: 7 were deletions, of which 5 were frameshift deletions and 2 were within introns. For the remaining 47 insertions, 23 were frameshift insertions, 5 were in-frame insertions ranging from 6-12 bp in length, and 19 were within introns (Figure 6). The proportion of indels in protein coding sequence that were in-frame was not significantly different from 1/3 (Fisher’s Exact Test p > 0.2). Again, these results suggest that selection did not substantially reduce the number of frameshift mutations captured during MA.

**Figure 6.**
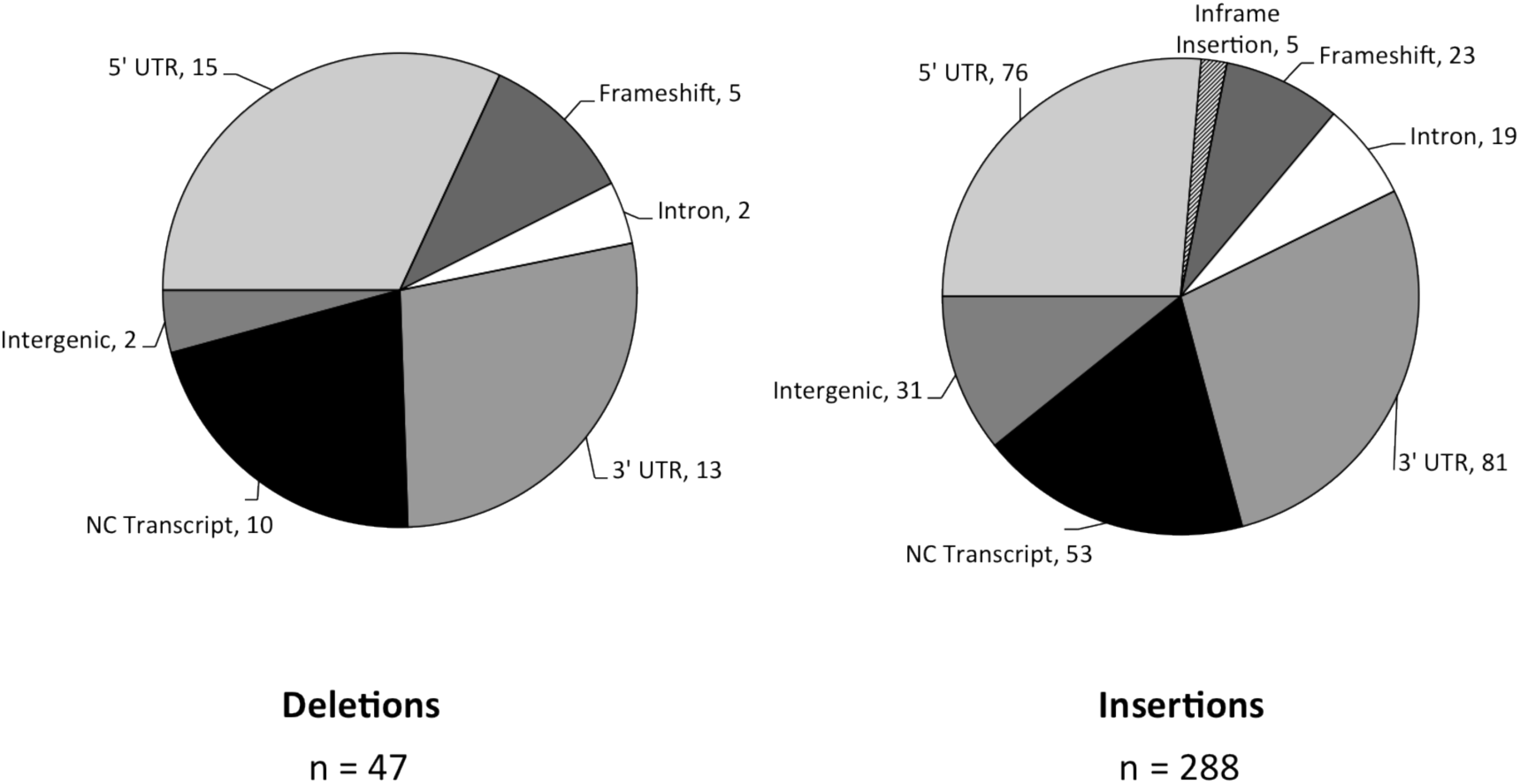
Summary of the locations and predicted functional consequences of the 335 small indels (less than 50bp in size). Numbers of indels identified across all 79 lines are shown for each category. Sum of the numbers in each pie chart is greater than the observed number of mutations because some indels had multiple predicted functional effects. NC = non-coding. UTR = transcribed, untranslated region.

#### Double Mutations, Complex Mutations and Structural Variants

As expected, no aneuploids were detected among the haploid MA lines. We specifically searched for other structural variants including medium-sized deletions (50-1000 bp), inversions, translocations and duplications using the DELLY software that predicts structural variants from short insert paired-end sequences (RAUSCH *et al.* 2012). We detected three medium-sized deletions, two of which were flanked by a number of SNMs (Supplementary Materials). In addition to multiple SNMs in the region of the three medium deletions, 16 other complex mutations and 17 separate double SNMs were also detected. Complex mutations thus occurred at a rate of 8.32 × 10^-12^ per base per generation and double SNMs occurred at a rate of 8.84 × 10^-12^ per base per generation.

#### Error in Identification of Mutations

Our *Sc. pombe* lines were haploids and our sequencing coverage exceeded 25x in every line (average = 44x). Given previous work in diploid *S. cerevisiae*, we predicted we would have a very low likelihood of incorrectly calling a SNM or indel. To confirm a low false positive error rate, we randomly chose five lines and checked all predicted mutations using Sanger sequencing (Supplementary Materials). Together, these lines contained 15 SNMs, 20 small insertions, 3 small deletions, 1 double SNM and 1 complex mutation. Sanger sequencing confirmed all of the mutations identified from next-generation sequencing, suggesting that the false positive error rate is likely is no larger than 0.025, and could be close to zero. We did find that in the region of the complex mutation (MA Line 58, Chromosome II, positions 4273519 – 4274319) additional mutations not identified in next-generation sequencing data were present (i.e. false negatives). These mutations were probably not detected due to the difficulty in reference mapping sequencing reads that are too divergent from the reference genome. This conjecture is supported by the extreme loss of coverage that was observed in this 800 bp region. Just 80 bp on either side of the region coverage was 48x, while across the complex mutation coverage averaged 9x, with 3 nucleotides in the center only being represented by a single read. Our results for this complex mutation suggest that the numbers of detected changes present in the 5 complex mutations we identified between the ancestor and reference strain and the 18 in the MA lines, are underestimates of the number actually present.

We also examined the probability of false negatives for SNMs. This was done by asking how many of the 272 mutations that differ between the reference and the ancestor were not found in each of the 79 MA lines. Any mutation present in the ancestor that is not identified in an MA line is a false negative, assuming the probability that a mutation converting the ancestor base back to the reference base at any of these sites is negligible. Given our estimate of the mutation rate, the probability of a mutation occurring at any of these sites in any line during the experiment is ∼0.007, indicating that failure to detect a site that differed in the ancestor is almost certainly a false negative. For SNMs, the false negative rate was 0.0001, while for small insertions and small deletions it was 0.011 and 0.017 respectively. Thus, we may have underestimated the base substitution, small insertion and small deletion mutation rates by 0.01, 1.1 and 1.7 percent respectively.

## DISCUSSION

### Rate of mutation at single nucleotides

We expected to observe a higher SNM rate in *Sc. pombe* than has been reported in *S. cerevisiae*, based on estimates of effective population size (SKELLY *et al.* 2009; BROWN *et al.* 2011) and on data using reporter genes (LYNCH 2010). Our estimate of the base pair mutation rate in *Sc. pombe*, based on the 326 accumulated SNMs, is 0.170 × 10^-9^ ± 0.013 per base per generation, which is almost identical to the mutation rate of 0.167 × 10^-9^ per base per generation, based on 867 accumulated SNM mutations in *S. cerevisiae* (ZHU *et al.* 2014). The effective population size in *Sc. pombe* was previously estimated as 1.0 × 10^7^, however this estimate uses the mutation rate estimate from haploid *S. cerevisiae* of 0.33×10^-9^ per base per generation (LYNCH *et al.* 2008) and the standing genetic variation among *Sc. pombe* strains (BROWN *et al.* 2011). With our estimate of the mutation rate, which is half as large, and the genome-wide estimate of π from Jefferies et al, 2015, we can recalculate the effective population size for *Sc. pombe* as 8.80 × 10^6^, essentially equal to our estimate of effective population size for *S. cerevisiae* 8.53 × 10^6^, considering its diploid genome (LITI *et al.* 2009). Thus, *Sc. pombe* and *S. cerevisiae* do not seem to differ in their effective population size, suggesting that selection should be similarly effective in both species.

#### Bias in single nucleotide mutations

The elevated rates of G/C → A/T mutations point to oxidative damage as a major role in mutagenesis in our MA lines (CHENG *et al.* 1992; SHEN *et al.* 1994). Within MA lines, G/C → A/T mutations occurred at a rate that was 2.98x greater than A/T → G/C mutations. The equilibrium G/C content calculated from the mutation spectra is 25% and 32% respectively (Figure S7). The difference between the two species is due to different mutational biases. In particular, G/C → A/T bias is 32% greater in *Sc. pombe* than in *S. cerevisiae*.

In both species the observed G/C content (35% in *Sc. pombe* and 38% in *S. cerevisiae*), is higher than predicted from mutation biases suggesting that there is either selection for lower G/C content or some other mechanism, perhaps biased gene conversion, which is causing the increase in genomic G/C content. Suggestive evidence for an additional force is present in the ancestor, whose G/C → A/T bias relative to the reference strain is 24% less than observed in the MA lines.

#### Rate and Spectrum of Indels

In our MA lines, we observed a strong insertion bias. Insertions accounted for 85% of small indel events, consistent with previous observations of the GT repeat region in the *ade6* gene of *Sc. pombe*, where 83% of indel events in wild-type were insertions (MANSOUR *et al.* 2001). A similar bias towards insertions has also been observed in *C. elegans* (3.75 insertions to deletions) (DENVER *et al.* 2004) and haploid *S. cerevisiae* (3.55 insertions to deletions) (LYNCH *et al.* 2008). In haploid *S. cerevisiae*, despite their rarity, the size of small deletions was able to negate the effect of small insertions in terms of changes in genome size. This is not the case in *Sc. pombe.* Even though small deletions were twice as large as small insertions, small insertions occurred six times more frequently resulting in a net gain of 340 bp across the genome. This suggests that *Sc. pombe*’s current genome size is not at equilibrium with respect to an insertion/deletion mutational balance. Additionally, in the ancestor small deletions may have been favored by selection. 17.3% of the differences between the ancestor and reference are small deletions compared to only 6.7% in the MA lines, which is a significant difference.

Similar to observations in *C. elegans, Sc. pombe* experiences small indels as often as SNMs (DENVER *et al.* 2004), resulting in an indel rate that is 10 fold higher than haploid *S. cerevisiae* and almost 35 fold higher than diploid *S. cerevisiae* (*Sc. pombe*: 0.174 × 10^-9^ vs. haploid *S. cerevisiae*: 0.2 × 10^-10^, and diploid *S. cerevisiae*: 0.5 × 10^-11^ per base pair per generation). One hypothesis for the differences in indel rate may be due to differences in genome complexity between the two species. *Sc. pombe*’s genome is 60.2% protein coding (57% excluding introns) while *S. cerevisiae*’s is 71% protein coding (70.5% excluding introns) indicating a higher prevalence of noncoding DNA in *Sc. pombe.* Most indels identified in *Sc. pombe* and *S. cerevisiae* are within low complexity, intergenic regions such as microsatellites and mononucleotide runs. However, in an analysis of short simple repetitive sequences, there is little observed difference in the amount of repetitive sequence in *S. cerevisiae* and *Sc. pombe* (KARAOGLU *et al.* 2005). An alternate hypothesis is that there are differences in mismatch repair pathways between these two species. However, both species contain MSH2 homologs, which are responsible for high fidelity repair of small indels (RUDOLPH *et al.* 1999), suggesting that this hypothesis would require differences in fidelity of the mismatch repair pathway, rather than presence versus absence. A third hypothesis is that there may be differences in polymerase fidelity between *Sc. pombe* and *S. cerevisiae* causing differing rates of strand slippage resulting in *Sc. pombe’s* increased insertion rate. However, given similar effective population sizes, we do not expect fidelity to differ in these two species.

#### Complex mutations, double mutations and aneuploidy

We also predicted an increase of mutations associated with double stranded breaks; particularly double SNMs and complex mutations. Double stranded breaks are lethal to a cell unless repaired. Repair can involve homologous recombination, which tends to be the preferred mechanism (RAJI and HARTSUIKER 2006), but can also utilize nonhomologous end-joining (NHEJ). Recombinational repair requires homologous copies of DNA, which in a haploid organism are present during the S and G2 phases of the cell cycle, when sister chromatids are present. In the absence of homologous DNA, double stranded breaks are repaired through NHEJ. Homologous recombination is considered an error-free method of repair, while non-homologous end joining is considered to be error-prone, introducing small insertions or deletions when partially degraded ends inhibit precise repair (CAVERO *et al.* 2007; SHRIVASTAV *et al.* 2008).

The locations of complex mutations were not random. Six of the 18 complex mutations occurred within three of the nine flocculin genes. All of these six mutations occurred within the characteristic tandem repeats found in these genes (VERSTREPEN *et al.* 2005). The flocculin tandem repeats are known to cause replication errors and are highly prone to double-stranded breaks and, as a result, recombination (VERSTREPEN *et al.* 2005). Their propensity for double stranded breaks could explain the repeated observation of complex mutations within them if complex mutations are, in fact, caused by mutagenic NHEJ repair of double-stranded breaks. To make sure that inferred mutations in the flocculin genes were not caused by gene conversion/recombination, we checked for and found no evidence of recombination between paralogs.

As we predicted given its haploid state, possession of only three chromosomes, and previous work (NIWA *et al.* 2006), there were no instances of aneuploidy. In *Sc. pombe* aneuploidy has only been observed as a disomic haploid of chromosome III. Even if a disomic haploid had been fixed in a line at an intermediate transfer, it would likely have been unstable (NIWA and YANAGIDA 1985; NIWA *et al.* 2006) and thus rapidly lost. Interestingly, when *S. cerevisiae* is passaged as a mitotic haploid it tends to be very unstable and at large population size, where selection is effective, it reverts to a diploid state (GLAZUNOV *et al.* 1988). This instability is in contrast to the relative stability of diploid strains (NISHANT *et al.* 2010; ZHU *et al.* 2014). This perhaps implies that while aneuploidy is more difficult in haploids than in diploids if all else is equal; when diploidy is the natural state, disomies, or other steps towards diploidy, may be tolerated and perhaps even favored in haploids. It would be interesting to see if the instability of haploidy in *S. cerevisiae*, which is a natural diploid, would also be reflected in instability of a diploid *Sc. pombe*, which is a natural haploid.

#### Cytosine mutation in absence of methylation

One of the major surprises of the *S. cerevisiae* mutation spectrum is the high mutation rate observed at some C:G base pairs, particularly in CpG dinucleotides found in CCG and TCG trinucleotides. We observed a similar, unexpectedly high mutation rate at some C:G base pairs, especially CCG and GCG. In many eukaryotes a major cause of mutation at cytosine nucleotides is spontaneous de-amination of methylated cytosines (5-mC) to thymine, which results in a T:G mismatch that can then be repaired, or replicated, to give a C:G → T:A substitution. Deamination of methylated cytosines was hypothesized to in part explain the elevated mutation rate of C:G base pairs in *S. cerevisiae* because there is some evidence for low levels of DNA methylation (TANG *et al.* 2012). *Sc. pombe* is not thought to possess methylation but, even if cytosines are methylated at a very low rate, methylation cannot explain the mutational bias at these sites. Methylated cytosines would cause C:G → T:A transition mutations, not the observed C:G to A:T transversions. Additionally, mutation biases at the C:G base pair in a CpG dinucleotide is no different from biases at a C:G base pair that is not present in a CpG. Together these observations suggest a different mechanism is driving the increased relative mutation rate at CpG dinucleotides other than deamination of methylated cytosines. This suggests that methylation may not be the only cause of high mutation rates at CpG dinucleotides in other species.

#### Spectrum of single nucleotide mutations

If differences between the ancestor and the reference represent both neutral and selected mutations, then the mutation spectrum might differ from that observed among the MA lines. In the MA lines SNMs represent 46.9% of the observed mutations, while in the ancestor they are significantly less common. The lower proportion of SNMs among differences between the ancestor and reference suggests that ∼1/3 of the SNMs that arose in the lineages linking the two strains were deleterious and were thus prevented from fixing. The location of the SNMs in the genome with respect to exon versus intron, and non-coding regions is similar in both the ancestor and MA lines suggesting that many of SNMs present in the ancestor relative to the reference are neutral (Figure 2, Figure S6).

#### Recently Published Work

A mutation accumulation study in *Sc. pombe* was recently published (FARLOW et al. 2015). While many of the results in that paper are in agreement with our findings, there are some striking differences. First, mutations in flocculation genes observed in that study were attributed to selection, whereas here they are attributed to genomic regions that are highly susceptible to double stranded breaks. Second, CpG mutations in that study were biased towards C:G → T:A, even though C:G → A:T mutations were more common averaged across all C:G sites. In contrast, we found C:G →A:T mutations were more common in CpG dinucleotides, and mutations at CpG sites did not vary from the genome wide mutation spectrum. Third, that study estimated the effective population size for *Sc. pombe* as ∼12 million, while we estimate the effective population size as ∼8.8 million. Lastly, the indel rate reported in that study is 3x smaller than the rate observed here.

The source of the conflicting results may be attributed to at least four possibilities: First, different *Sc. pombe* strains were used in each experiment. We used *Sc. pombe* 972h-, which is the standard lab strain and the basis of the *Sc. pombe* reference genome (WOOD *et al.* 2002). The other study used a strain from the genome-wide deletion mutant library produced by BIONEER derived from *Sc. pombe* 975h+. While both strains were originally isolated from the same population (LEUPOLD 1950), genetic differences may have arisen that could account for these differences.

Second, the culture conditions vary between the two experiments; our conditions (incubation at 30°C for 48h on YPD agar plates) were chosen in order to be identical to the conditions used in the *S. cerevisiae* MA study (ZHU *et al.* 2014) and also happened to be the optimal growth temperature for our isolate (http://www.atcc.org/Products/All/26189.aspx). The other study cultured their MA lines at 32°C on 3-4 day transfer cycles.

Third, the methods of sequence mapping and variant calling differ between the two studies. If there is no substantial difference in mutation rate and spectrum between the two strains, the fact that we identify more indels suggests that our methods reliably identify more of the mutations (i.e. we have fewer false negatives). We estimated our false negative error rate for SNMs and indels and, in all cases, they are less than 2%. A direct estimate of false negative error rate was not performed in the other study, though they did estimate a false negative rate by adding simulated mutations into their reference assemblies and attempting to detect them.

Finally, there appears to be at least one error in the calculations of parameters. Using their estimate of π (given as 0.003 in their Table S3) and their genome-wide single-nucleotide mutation rate (1.7 × 10^-10^ per base per generation), we calculate an effective population size of ∼7.5 million. This corrected estimate using their data is very similar to our estimate of ∼8.8 million. We have been unable to replicate their estimate of 12 million using their data.

In summary, we find that in contrast to previous findings, *Sc. pombe* and *S. cerevisiae* have essentially identical base pair mutation rates, which, coupled with previous estimates of within-species polymorphisms, suggests their effective population sizes are essentially identical. We observe considerable differences in the mutation spectrum and indel rate suggesting that there are species-specific differences in factors affecting mutation bias and indel rate between these two species. Additionally, we note that small insertions versus deletions predicts a growing genome, suggesting either that the genome size in *Sc. pombe* is not at equilibrium, or that rare large deletions offset increases caused by small indel bias. The sample size of captured large insertions and deletions was insufficient to test this hypothesis. Finally, we found that CpG sites are highly mutagenic, but the mutation bias at these sites is not caused by deamination of methylated cytosine, suggesting another factor makes these sites prone to mutation.

## References

Andolfatto, P., 2005 Adaptive evolution of non-coding DNA in *Drosophila*. Nature 437: 1149–1152.

Antequera, F., M. Tamame, J. Villanueva and T. Santos, 1984 DNA methylation in the fungi. Journal of Biological Chemistry 259: 8033–8036.

Aronesty, E., 2011 ea-utils: Command-line tools for processing biological sequencing data, pp.

Bell, J., 2008 A simple way to treat PCR products prior to sequencing using ExoSAP-IT. Biotechniques 44: 834.

Bestor, T. H., and G. L. Verdine, 1994 DNA methyltransferases. Current opinion in cell biology 6: 380–389.

Blumenstiel, J. P., A. C. Noll, J. A. Griffiths, A. G. Perera, K. N. Walton et al., 2009 Identification of EMS-induced mutations in *Drosophila melanogaster* by whole-genome sequencing. Genetics 182: 25–32.

Brown, W. R., G. Liti, C. Rosa, S. James, I. Roberts et al., 2011 A geographically diverse collection of *Schizosaccharomyces pombe* isolates shows limited phenotypic variation but extensive karyotypic diversity. G3: Genes, Genomes, Genetics 1: 615–626.

Cavero, S., C. Chahwan and P. Russell, 2007 Xlf1 is required for DNA repair by nonhomologous end joining in *Schizosaccharomyces pombe*. Genetics 175: 963–967.

Cheng, K. C., D. S. Cahill, H. Kasai, S. Nishimura and L. A. Loeb, 1992 8-Hydroxyguanine, an abundant form of oxidative DNA damage, causes G-T and A-C substitutions. Journal of Biological Chemistry 267: 166–172.

Daley, J. M., P. L. Palmbos, D. Wu and T. E. Wilson, 2005 Nonhomologous end joining in yeast. Annu. Rev. Genet. 39: 431–451.

Danecek, P., A. Auton, G. Abecasis, C. A. Albers, E. Banks et al., 2011 The variant call format and VCFtools. Bioinformatics 27: 2156–2158.

Denver, D. R., S. Feinberg, C. Steding, M. Durbin and M. Lynch, 2006 The relative roles of three DNA repair pathways in preventing *Caenorhabditis elegans* mutation accumulation. Genetics 174: 57–65.

Denver, D. R., K. Morris, M. Lynch and W. K. Thomas, 2004 High mutation rate and predominance of insertions in the *Caenorhabditis elegans* nuclear genome. Nature 430: 679–682.

Denver, D. R., L. J. Wilhelm, D. K. Howe, K. Gafner, P. C. Dolan et al., 2012 Variation in base-substitution mutation in experimental and natural lineages of *Caenorhabditis* nematodes. Genome biology and evolution: evs028.

Douzery, E. J., E. A. Snell, E. Bapteste, F. Delsuc and H. Philippe, 2004 The timing of eukaryotic evolution: does a relaxed molecular clock reconcile proteins and fossils? Proceedings of the National Academy of Sciences of the United States of America 101: 15386–15391.

Farlow, A., H. Long, S. Arnoux, W. Sung, T. G. Doak et al., 2015 The Spontaneous Mutation Rate in the Fission Yeast *Schizosaccharomyces pombe*. Genetics: genetics. 115.177329.

Glazunov, A., A. Boreiko and A. Esser, 1988 [Relative competitiveness of haploid and diploid yeast cells growing in a mixed population]. Mikrobiologiia 58: 769–777.

Gordon, A., and G. Hannon, 2010 Fastx-toolkit. FASTQ/A short-reads preprocessing tools (unpublished) http://hannonlab.cshl.edu/fastx_toolkit.

Graur, D., 2003 Single-base Mutation. Nature encyclopedia of the human genome: 287.

Greene, E. A., C. A. Codomo, N. E. Taylor, J. G. Henikoff, B. J. Till et al., 2003 Spectrum of chemically induced mutations from a large-scale reverse-genetic screen in *Arabidopsis*. Genetics 164: 731–740.

Hall, D. W., R. Mahmoudizad, A. W. Hurd and S. B. Joseph, 2008 Spontaneous mutations in diploid *Saccharomyces cerevisiae*: another thousand cell generations. Genetics research 90: 229–241.

Halligan, D. L., and P. D. Keightley, 2009 Spontaneous mutation accumulation studies in evolutionary genetics. Annual Review of Ecology, Evolution, and Systematics 40: 151–172.

Heckman, D. S., D. M. Geiser, B. R. Eidell, R. L. Stauffer, N. L. Kardos et al., 2001 Molecular evidence for the early colonization of land by fungi and plants. Science 293: 1129–1133.

Heichinger, C., C. J. Penkett, J. Bähler and P. Nurse, 2006 Genome-wide characterization of fission yeast DNA replication origins. The EMBO Journal 25: 5171–5179.

Hershberg, R., and D. A. Petrov, 2008 Selection on codon bias. Annual review of genetics 42: 287–299.

Ho, S. Y., M. J. Phillips, A. Cooper and A. J. Drummond, 2005 Time dependency of molecular rate estimates and systematic overestimation of recent divergence times. Molecular Biology and evolution 22: 1561–1568.

Hoffman, P. D., J. M. Leonard, G. E. Lindberg, S. R. Bollmann and J. B. Hays, 2004 Rapid accumulation of mutations during seed-to-seed propagation of mismatch-repair-defective *Arabidopsis*. Genes & development 18: 2676–2685.

Holbeck, S. L., and J. N. Strathern, 1997 A role for REV3 in mutagenesis during double-strand break repair in *Saccharomyces cerevisiae*. Genetics 147: 1017–1024.

Humphrey, T., 2000 DNA damage and cell cycle control in *Schizosaccharomyces pombe*. Mutation Research/Fundamental and Molecular Mechanisms of Mutagenesis 451: 211–226.

Jeffares, D. C., C. Rallis, A. Rieux, D. Speed, M. Převorovský et al., 2015 The genomic and phenotypic diversity of *Schizosaccharomyces pombe*. Nature genetics 47: 235–241.

Joseph, S. B., and D. W. Hall, 2004 Spontaneous mutations in diploid Saccharomyces cerevisiae more beneficial than expected. Genetics 168: 1817–1825.

Karaoglu, H., C. M. Y. Lee and W. Meyer, 2005 Survey of simple sequence repeats in completed fungal genomes. Molecular Biology and evolution 22: 639–649.

Keightley, P. D., U. Trivedi, M. Thomson, F. Oliver, S. Kumar et al., 2009 Analysis of the genome sequences of three *Drosophila melanogaster* spontaneous mutation accumulation lines. Genome research: gr. 091231.091109.

Koornneeff, M., L. Dellaert and J. Van der Veen, 1982 EMS-and relation-induced mutation frequencies at individual loci in *Arabidopsis thaliana (L.) Heynh*. Mutation Research/Fundamental and Molecular Mechanisms of Mutagenesis 93: 109–123.

Leupold, U., 1950 Die vererbung von homothallie und heterothallie bei Schizosaccharomyces pombe. CR Trav. Lab. Carlsberg Ser. Physiol. 24: 381–480.

Levinson, G., and G. A. Gutman, 1987 Slipped-strand mispairing: a major mechanism for DNA sequence evolution. Molecular Biology and evolution 4: 203–221.

Li, H., and R. Durbin, 2009 Fast and accurate short read alignment with Burrows–Wheeler transform. Bioinformatics 25: 1754–1760.

Liti, G., D. M. Carter, A. M. Moses, J. Warringer, L. Parts et al., 2009 Population genomics of domestic and wild yeasts. Nature 458: 337–341.

Lynch, M., 2010 Evolution of the mutation rate. Trends in Genetics 26: 345–352.

Lynch, M., W. Sung, K. Morris, N. Coffey, C. R. Landry et al., 2008 A genome-wide view of the spectrum of spontaneous mutations in yeast. Proceedings of the National Academy of Sciences 105: 9272–9277.

Mansour, A. A., C. Tornier, E. Lehmann, M. Darmon and O. Fleck, 2001 Control of GT repeat stability in *Schizosaccharomyces pombe* by mismatch repair factors. Genetics 158: 77–85.

McKenna, A., M. Hanna, E. Banks, A. Sivachenko, K. Cibulskis et al., 2010 The Genome Analysis Toolkit: a MapReduce framework for analyzing next-generation DNA sequencing data. Genome research 20: 1297–1303.

McLaren, W., B. Pritchard, D. Rios, Y. Chen, P. Flicek et al., 2010 Deriving the consequences of genomic variants with the Ensembl API and SNP Effect Predictor. Bioinformatics 26: 2069–2070.

Muller, H. J., 1930 Types of visible variations induced by X-rays in *Drosophila*. Journal of Genetics 22: 299–334.

Nachman, M. W., and S. L. Crowell, 2000 Estimate of the mutation rate per nucleotide in humans. Genetics 156: 297–304.

Ness, R. W., A. D. Morgan, N. Colegrave and P. D. Keightley, 2012 Estimate of the spontaneous mutation rate in *Chlamydomonas reinhardtii*. Genetics 192: 1447–1454.

Nishant, K., W. Wei, E. Mancera, J. L. Argueso, A. Schlattl et al., 2010 The baker's yeast diploid genome is remarkably stable in vegetative growth and meiosis. PLOS Genetics 6: e1001109.

Niwa, O., Y. Tange and A. Kurabayashi, 2006 Growth arrest and chromosome instability in aneuploid yeast. Yeast 23: 937–950.

Niwa, O., and M. Yanagida, 1985 Triploid meiosis and aneuploidy in *Schizosaccharomyces pombe*: an unstable aneuploid disomic for chromosome III. Current genetics 9: 463–470.

Raji, H., and E. Hartsuiker, 2006 Double-strand break repair and homologous recombination in *Schizosaccharomyces pombe*. Yeast 23: 963–976.

Rausch, T., T. Zichner, A. Schlattl, A. M. Stütz, V. Benes et al., 2012 DELLY: structural variant discovery by integrated paired-end and split-read analysis. Bioinformatics 28: i333–i339.

Robinson, J. T., H. Thorvaldsdóttir, W. Winckler, M. Guttman, E. S. Lander et al., 2011 Integrative genomics viewer. Nature biotechnology 29: 24–26.

Rozen, S., and H. Skaletsky, 1999 Primer3 on the WWW for general users and for biologist programmers, pp. 365–386 in Bioinformatics methods and protocols. Springer.

Rudolph, C., C. Kunz, S. Parisi, E. Lehmann, E. Hartsuiker et al., 1999 The *msh2* Gene of *Schizosaccharomyces pombe* Is Involved in Mismatch Repair, Mating-Type Switching, and Meiotic Chromosome Organization. Molecular and cellular biology 19: 241–250.

Rutter, M. T., A. Roles, J. K. Conner, R. G. Shaw, F. H. Shaw et al., 2012 Fitness of *Arabidopsis thaliana* mutation accumulation lines whose spontaneous mutations are known. Evolution 66: 2335–2339.

Saxer, G., P. Havlak, S. A. Fox, M. A. Quance, S. Gupta et al., 2012 Whole genome sequencing of mutation accumulation lines reveals a low mutation rate in the social amoeba *Dictyostelium discoideum*. PLoS One 7: e46759.

Schneider, C. A., W. S. Rasband and K. W. Eliceiri, 2012 NIH Image to ImageJ: 25 years of image analysis. Nature methods 9: 671–675.

Shen, J.-C., W. M. Rideout and P. A. Jones, 1994 The rate of hydrolytic deamination of 5-methylcytosine in double-stranded DNA. Nucleic acids research 22: 972–976.

Shrivastav, M., L. P. De Haro and J. A. Nickoloff, 2008 Regulation of DNA double-strand break repair pathway choice. Cell research 18: 134–147.

Skelly, D. A., J. Ronald, C. F. Connelly and J. M. Akey, 2009 Population genomics of intron splicing in 38 *Saccharomyces cerevisiae* genome sequences. Genome biology and evolution 1: 466–478.

Strathern, J. N., B. K. Shafer and C. B. McGill, 1995 DNA synthesis errors associated with double-strand-break repair. Genetics 140: 965–972.

Sung, W., M. S. Ackerman, S. F. Miller, T. G. Doak and M. Lynch, 2012a Drift-barrier hypothesis and mutation-rate evolution. Proceedings of the National Academy of Sciences 109: 18488–18492.

Sung, W., A. E. Tucker, T. G. Doak, E. Choi, W. K. Thomas et al., 2012b Extraordinary genome stability in the ciliate *Paramecium tetraurelia*. Proceedings of the National Academy of Sciences 109: 19339–19344.

Tang, Y., X.-D. Gao, Y. Wang, B.-F. Yuan and Y.-Q. Feng, 2012 Widespread existence of cytosine methylation in yeast DNA measured by gas chromatography/mass spectrometry. Analytical chemistry 84: 7249–7255.

Trapnell, C., L. Pachter and S. L. Salzberg, 2009 TopHat: discovering splice junctions with RNA-Seq. Bioinformatics 25: 1105–1111.

Trapnell, C., B. A. Williams, G. Pertea, A. Mortazavi, G. Kwan et al., 2010 Transcript assembly and quantification by RNA-Seq reveals unannotated transcripts and isoform switching during cell differentiation. Nature biotechnology 28: 511–515.

Verstrepen, K. J., A. Jansen, F. Lewitter and G. R. Fink, 2005 Intragenic tandem repeats generate functional variability. Nature genetics 37: 986–990.

Wood, V., R. Gwilliam, M.-A. Rajandream, M. Lyne, R. Lyne et al., 2002 The genome sequence of *Schizosaccharomyces pombe*. Nature 415: 871–880.

Wood, V., M. A. Harris, M. D. McDowall, K. Rutherford, B. W. Vaughan et al., 2011 PomBase: a comprehensive online resource for fission yeast. Nucleic acids research: gkr853.

Zhu, Y. O., M. L. Siegal, D. W. Hall and D. A. Petrov, 2014 Precise estimates of mutation rate and spectrum in yeast. Proceedings of the National Academy of Sciences 111: E2310–E2318.

